# Spatial organization and function of RNA molecules within phase-separated condensates are controlled by Dnd1

**DOI:** 10.1101/2023.07.09.548244

**Authors:** Kim Joana Westerich, Katsiaryna Tarbashevich, Jan Schick, Antra Gupta, Mingzhao Zhu, Kenneth Hull, Daniel Romo, Dagmar Zeuschner, Mohammad Goudarzi, Theresa Gross-Thebing, Erez Raz

**Affiliations:** Institute of Cell Biology, Center for Molecular Biology of Inflammation, University of Münster; 48149 Münster, Germany; Department of Chemistry & Biochemistry and The Baylor Synthesis and Drug-Lead Discovery Laboratory, Baylor University, Waco, Texas 76706, United States; Electron Microscopy Facility, Max Planck Institute for Molecular Biomedicine, 48149 Münster, Germany; Max Planck Institute for Molecular Biomedicine, 48149 Münster, Germany

**Keywords:** Zebrafish, germ cells, Dead end, *nanos*, RNA translation, germ granules, phase separation

## Abstract

Germ granules, condensates of phase-separated RNA and protein, are organelles essential for germline development in different organisms The patterning of the granules and its relevance for germ cell fate are not fully understood. Combining three-dimensional *in vivo* structural and functional analyses, we study the dynamic spatial organization of molecules within zebrafish germ granules. We find that localization of RNA molecules to the periphery of the granules, where ribosomes are localized depends on translational activity at this location. In addition, we find that the vertebrate-specific Dead end (Dnd1) protein is essential for *nanos3* RNA localization at the condensates’ periphery. Accordingly, in the absence of Dnd1, or when translation is inhibited, *nanos3* RNA translocates into the granule interior, away from the ribosomes, a process that is correlated with loss of germ cell fate. These findings highlight the relevance of sub-granule compartmentalization for posttranscriptional control, and its importance for preserving germ cell totipotency.

## INTRODUCTION

Germline development relies on conserved RNA and protein molecules that are highly enriched within germ granules, phase-separated, non-membrane-bound condensates^1–6^. The RNAs that reside within these organelles encode for proteins that are required during different phases of germline development, demanding a precise regulation over their translation^7, 8^. Indeed, many of the proteins located within germ granules were shown to have RNA-binding properties, consistent with the idea that the granules are organelles where post-transcriptional regulation occurs^7, 9^. Recent studies suggest that germ granules serve as protective storage sites for repressed mRNAs^10, 11^, a role that could be especially critical in the case of maternally-provided RNAs, whose translation is required at different time points of germ cell development^3, 7, 8, 12^. Interestingly, while many germline determinants reside within the granules, at least in some cases and stages of development, their actual integration into phase-separated structures is not essential for their function^10^. Together, despite their importance for the development of germ cells, the mechanisms that control the spatial organization of molecules within the condensates and the relevance of their organization for the function of the organelle are not fully understood^3–5^.

An attractive model for studying germ granule organization and function in vertebrates is zebrafish, in which the germline condensates are relatively large and accessible for *in vivo* imaging^13^. A key component localized to these organelles is Dead end (Dnd1), a conserved vertebrate-specific RNA-binding protein that is essential for germline development. Loss of the protein can result in sterility and formation of germ cell tumors^14–20^ and accordingly, recent work showed that germ cells with impaired Dnd1 function fail to maintain their totipotency and acquire somatic fates^21^. Despite the central role of Dnd1 in controlling the development of the vertebrate germline, the precise molecular function of the protein in the context of the phase-separated structures within which it resides is unknown.

In zebrafish, Dnd1 was shown to interact with the mRNA that encodes for the germ granule-localized protein Nanos3^22^. In this case, Dnd1 was shown to counteract miRNAs that would otherwise inhibit *nanos3* RNA function in the germ cells^22^. The function of Nanos is in turn essential for germ cell development in different vertebrate and invertebrate organisms^23–27^, and localization of its RNA to the germ plasm has been linked to its translation state in *Drosophila*^28^. Here, we reveal a role for Dnd1 in regulating the spatial distribution of *nanos3* RNA within germ granules, which facilitates the translation of the RNA, thereby ensuring proper development of the germline.

## RESULTS

### Nanos3 preserves germ cell fate and its mRNA is spatially associated with Dnd1 protein clusters in germ granules

Our recent finding that germ cells depleted of Dnd1 adopt somatic cell fates^21^ prompted us to determine whether inhibiting Nanos3 could affect the cells in a similar way. To investigate this, we employed embryos carrying a mutation in the gene encoding for the chemokine Cxcl12a that prevents germ cells from migrating toward the gonad, such that they reside at ectopic positions throughout the embryo (Figure 1A)^29^. Interestingly, while non-manipulated ectopic germ cells maintain their morphology independent of the tissue in which they reside, germ cells depleted of Nanos3 protein adopt various somatic-like cell shapes (e.g., that of a muscle cell, (Figure 1A, 1B)), consistent with the idea that Dnd1 and Nanos3 function in the same pathway required for maintaining germ cell fate^21^.

**Figure 1.**
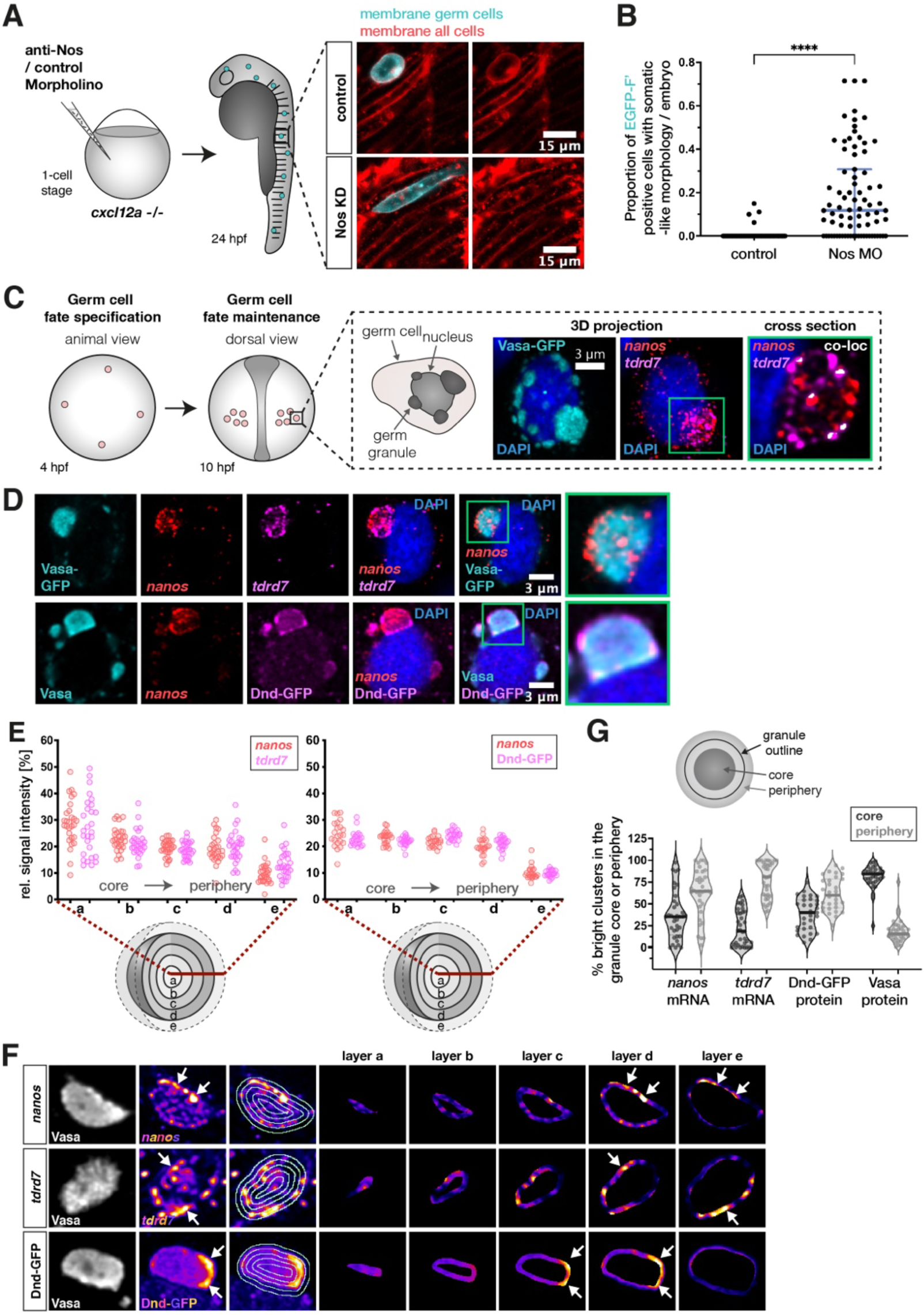
Nanos3 maintains germ cell fate, and its mRNA assembles in distinct clusters within germ granules. (A) 1-cell stage embryos that lack the chemokine guidance cue Cxcl12a were injected with a morpholino antisense oligonucleotide (anti-Nos Morpholino), inhibiting the translation of nanos3 mRNA. The germ cells were then evaluated for somatic cell morphologies at 24 hpf. Images show an example of a Nanos3-deficient germ cell exhibiting elongated muscle precursor-like appearance. (B) Evaluation of the proportion of wildtype and Nanos3-deficient germ cells with soma-like morphologies. Cells were only counted as soma-like when they displayed clear morphologies of neurons, muscle or notochord cells. (C) Scheme depicting early germ cell development. Germ cells are specified at 4 hpf, become migratory and form two bilateral clusters at 10 hpf. The black box shows a representative image of perinuclear germ granules marked by the Vasa-GFP protein, with enrichment of the endogenous germline mRNAs tdrd7 and nanos3 (detected by RNAscope probes). (D) Distribution of nanos3 mRNA relative to tdrd7 mRNA (upper panels) and Dnd1 protein (lower panels) within germ granules marked by Vasa-GFP (upper panels) or endogenous Vasa protein (lower panels). One confocal plane is presented. (E) 3D analysis of the distribution of germ granule components across concentric layers of the condensate. (F) Heat map images of germ granule single confocal planes showing the distribution of nanos3 mRNA, tdrd7 mRNA or Dnd1 protein in different layers of the condensate. White arrows point at distinct clusters of these molecules, primarily detected in peripheral layers (layer d, e) of the condensate. (G) Proportion of bright clusters of nanos3 mRNA, tdrd7 mRNA, Dnd-GFP protein or Vasa protein detected in the core (dark grey) or peripheral (light grey) regions of the granule. The area occupied by bright clusters in the core or periphery was normalized to the size of the respective region. The peripheral region also includes the immediate area outside of the granule. Each dot in the graph represents the value derived from a single germ granule. For each granule, the brightest 5% of pixels were defined as bright clusters. Single confocal planes were used for the analysis. n (nanos, tdrd7) = 36, n (Vasa) = 35, n (Dnd-GFP) = 31. N = 3. See also Figure S1.

As a first step in investigating the function of the Dnd1 protein within germ granules^30^, we examined if the condensates are spatially organized with respect to the distribution of molecules, particularly *nanos3* mRNA, and if so, whether Dnd1 plays a role in controlling it. Here, we took advantage of the fact that during early stages of zebrafish embryonic development (up to 10 hours post fertilization (hpf)), germ granules can be relatively large, reaching sizes of over 50 µm^3^ (Figures S1A and S2A), thus allowing for high spatial resolution analyses *in vivo*. To this end, we employed the conserved germ granule protein Vasa^13, 31, 32^ for determining the volume and shape of the organelle (Figures 1C and S1B, Movie S1). Focusing on the organization of germline mRNAs, we evaluated the distribution of *nanos3* and *tudor domain containing 7* (*tdrd7*) mRNAs within the germ granules. Both mRNAs showed enrichment within the condensates in largely non-overlapping punctate sub-granule clusters (Figures 1C and S1A). The distinct distribution pattern of *nanos3* and *tdrd7* RNA molecules (Figure S1A, Movie S1) is similar to homotypic RNA clusters that were first observed in *Drosophila* germ granules^33–36^.

To characterize the distribution of different mRNAs and proteins within the phase-separated condensates, we developed a 3D signal-visualization approach that allowed us to determine the spatial organization of molecules along the core-to-periphery axis of the organelles (Figure S1B). We selected condensates with a minimum radius of 0.8 µm to obtain a sufficient sub-granule resolution. Interestingly, our analysis revealed that *nanos3* mRNA, *tdrd7* mRNA and the Dnd1 protein are found throughout the condensate, including domains that extend beyond the organelle’s border as defined by Vasa (Figures 1D and 1E, Movie S1). Although germ granules vary in size, we found that within the size range we focused on (radius = 0.8-2.5 µm), the spatial distribution of the mRNAs is similar (Figure S2C).

While we detected Dnd1 throughout the granule, in contrast to Vasa protein, Dnd1 formed distinct discontinuous clusters within the condensates, often located in vicinity of *nanos3* RNA clusters (Figure 1D, lower panels). Intriguingly, we found that such clusters of enriched protein or RNA material are mostly localized to the periphery of the organelle (Figures 1F and 1G). Thus, although different germline components such as *nanos3*, *tdrd7* and Dnd1 are found throughout the germ granule, they are preferentially enriched in distinct locations at the condensate periphery.

As Dnd1 was shown to interact with *nanos3* mRNA^22^, we next set out to examine the spatial relationship between the two molecules within the germ granules. Interestingly we found that Dnd1 protein and *nanos3* mRNA are localized to adjacent, partially overlapping clusters (Figures S1C, 2A and 2B). To address whether the adjacent clusters could be associated with one another, we followed the localization of these molecules in live embryos employing the PP7 system^37, 38^ for detection of *nanos3* RNA. Indeed, while Dnd1 protein and *nanos3* mRNA clusters were highly dynamic, they maintained their proximity within the condensate over time (Figure 2C, Movie S2).

**Figure 2.**
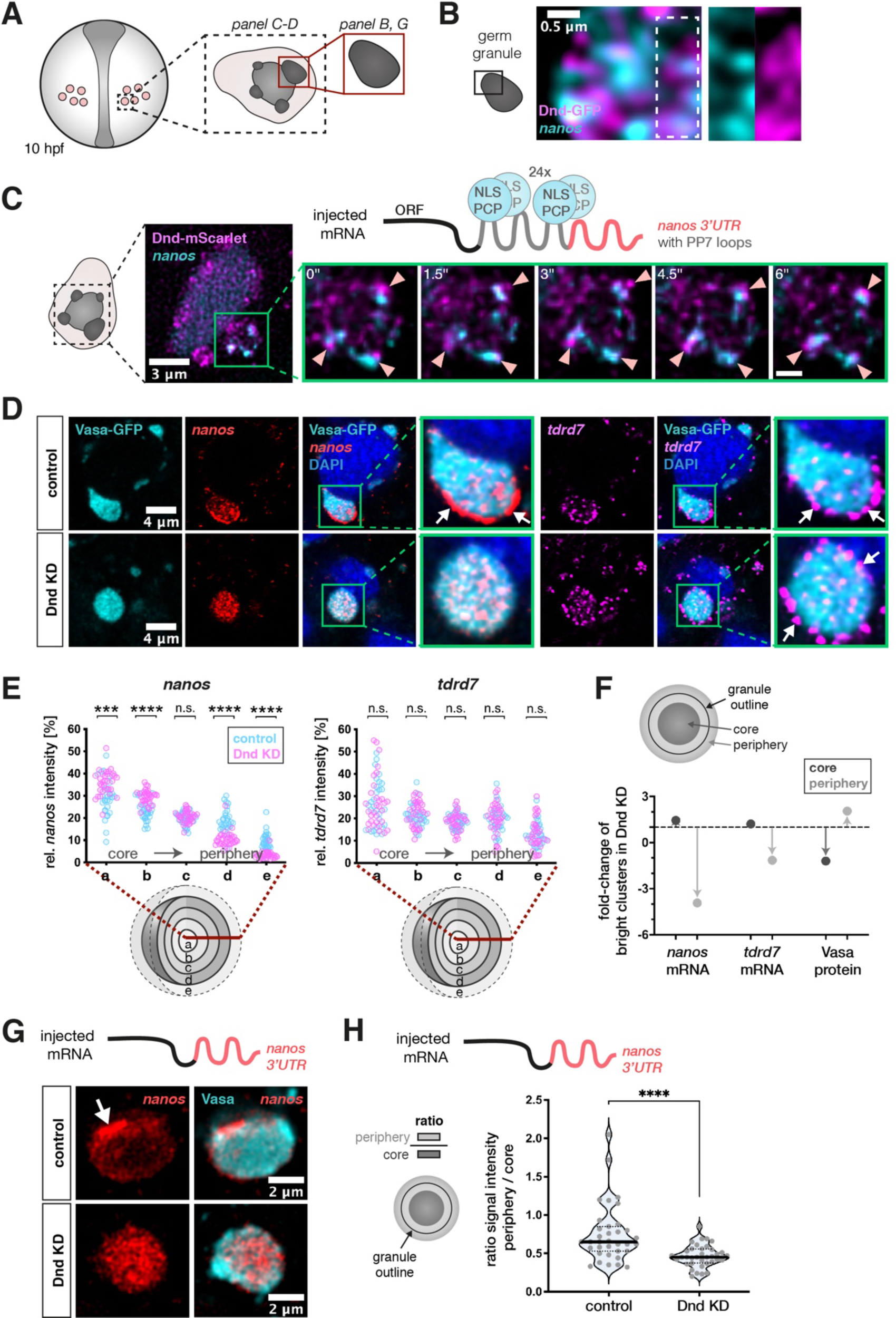
Dnd1 is required for maintaining *nanos3* mRNA clusters at the periphery of germ granule condensates. (A) A scheme showing germ cells in a 10 hpf embryo. The black dotted box, and the red box depict the region of interest for the indicated image panels. (B) Localization of Dnd-GFP protein clusters and endogenous *nanos3* mRNA within the germ granule. White dotted box highlights the magnified region shown in separate channels in the right panels. (C) The mobility of protein and mRNA clusters within germ granules in embryos injected with mRNA containing the *nanos3* open reading frame (ORF) and PP7 stem loops (grey in the scheme) inserted in the *nos 3’UTR* (red in the scheme), along with mRNA encoding for the PP7 loop-detecting NLS-PCP-YFP coat protein (light blue) and mRNA encoding for the Dnd-mScarlet protein (magenta). Green box shows a single large germ granule with *nanos3* mRNA and Dnd1 protein clusters. Arrowheads in the magnification panels point at mRNA and protein clusters within this granule that remain in proximity over time. Time interval = 1.5 seconds, scale bar = 0.8 µm (see Movie S1). Images are shown as maximum intensity projections comprising 4 focal planes with a z-distance of 500 nm. (D) Distribution of endogenous *nanos3* and *tdrd7* mRNA within germ granules (marked by Vasa-GFP) in control and Dnd knockdown (KD) embryos. Green boxes show magnifications of large germ granule condensates from the image panels, with arrows pointing at mRNA enrichment at the granule periphery. Single-plane images are presented. (E) 3D analysis of the distribution of endogenous *nanos3* and *tdrd7* mRNAs across germ granule concentric layers under control and Dnd KD conditions. Statistical significance was determined using Mann-Whitney *U* test. ***P = 0.0002, ****P< 0.0001. n (control) = 28, n (Dnd KD) = 32. N = 3. (F) Shown is the effect of Dnd knockdown on the proportion of bright clusters of *nanos3* mRNA, *tdrd7* mRNA or Vasa protein in the granule core (dark grey) or periphery (light grey). Arrows indicate the increase or decrease of the proportion of clusters relative to wildtype (indicated by the dotted line). The area occupied by bright clusters in the core or periphery was normalized to the size of the respective region. The peripheral region also includes the immediate area outside of the granule. For each granule, the brightest 5% of pixels were defined as bright clusters. Single confocal planes were used for the analysis. n (control) = 31, n (Dnd KD) = 32. N = 3. (G) Single-plane confocal image presenting the localization of injected *nos 3’UTR*-containing mRNA within germ granules visualized by endogenous Vasa protein under control and Dnd KD conditions. (H) A graph showing the ratio of the *nanos3’UTR*-contaning RNA level between the granule periphery and the core of the condensate. Significance was determined using Mann-Whitney *U* test. ****P<0.0001. n (control) = 35, n (Dnd KD) = 36. N = 3. See also Figures S1 and S2.

Together, our findings demonstrate that zebrafish germ granules are patterned organelles, in which RNAs and proteins occupy distinct domains that maintain their spatial relationships, as exemplified by *nanos3* mRNA and Dnd1 protein. This suggests cross dependency in localization among molecules and functional significance for sub-granule organization.

### Dnd1 is required for maintaining *nanos3* mRNA clusters at the periphery of germ granule condensates

We have previously shown that *nanos3* mRNA is a target of Dnd1^22^, that knocking down Dnd1 results in loss of germ cell pluripotency^21^ and show here that knocking down Nanos3 results in a similar phenotype, where germ cells differentiate into somatic cells (Figure 1A). These findings, together with the spatial relationships we identified between Dnd1 protein and *nanos3* RNA, prompted us to ask whether Dnd1 controls *nanos3* distribution within the granule. To investigate this, we examined the organization of germ granules in 10 hpf live embryos following morpholino-mediated inhibition of *dnd1* RNA translation (Figures 2D and 2E). Although Dnd1 loss eventually leads to a decrease in germ granule size and number at later stages of development^21^, we found that, at least until 10 hpf, the volume of large germ granules is not altered (Figure S2A). However, we detected a pronounced effect on the distribution of *nanos3* mRNA within the granules, as evaluated by the 3D granule layer analysis. In particular, Dnd1 depletion caused a strong reduction in *nanos3* levels at the periphery of the condensate, concomitant with an increase in the mRNA level at the core (Figure 2D and 2E). Consistently, the number of bright clusters of *nanos3* RNA in the granule periphery was strongly reduced in cells knocked down for Dnd1 (Figure 2F). This effect was mediated by the 3’UTR of *nanos3* (*nos 3’UTR*), since injected RNA that contained this Dnd1-binding element^22^ was depleted from the periphery of the granule as well (Figure 2G and 2H). Critically, these results do not represent a global effect on RNA distribution, since the localization of *tdrd7* mRNA was not altered under these conditions (Figures 2D and 2E). Thus, the distribution of different RNA molecules within germ granules can be regulated by distinct molecular mechanisms. Interestingly however, despite the dramatic effect of Dnd1 knockdown on the positioning of *nanos3* mRNA clusters within the granules, the discrete homotypic clustering of *tdrd7* and *nanos3* molecules was preserved (Figure S2B). These findings show that the basis for segregating RNA species from one another differs from the mechanisms that control the spatial distribution of the clusters within the granule.

### Zebrafish germ granule condensates contain translation-repression factors and exhibit translation activity and enrichment of ribosomes at their periphery

The altered distribution of *nanos3* mRNA within the germ granule observed upon Dnd1 knockdown led us to examine whether the change in the spatial organization of the mRNA could have a functional significance. As a first step in this analysis, we sought to determine the localization of factors that could regulate mRNA function. In line with recent findings in invertebrates suggesting that translationally silent mRNA molecules are stored within phases-separated granules^10, 11^, we could detect the RNA-induced silencing complex (RISC) components *mir-430* and Argonaute protein 2 (Ago2) within zebrafish germ granules (Figures 3A and 3B). In addition, we found the translation initiation factors eIF4G, eIF4E and PolyA-binding protein (PABP) within the organelles (Figure S3F). Intriguingly however, while the translation initiation factors were distributed throughout the granule (Figure S3F), we found that ribosomes are primarily localized to the edge of the condensate, as detected by antibodies targeting large (L10a) and small (S2 and S6) ribosomal subunit proteins (Figures 3C and S3E). These observations were backed up by electron microscope (EM) data showing distinct enrichment of ribosomes at the germ granule periphery, and presence of polyribosomes at this location (Figures 3D and 3E). Glycogen particles that were previously detected in germ cells^39–42^, were also found in the granules, where they could be well distinguished from ribosomes by their size and staining intensity (Figure S3D). Interestingly, the EM images also revealed the presence of rough endoplasmatic reticulum (ER) membrane at the circumference of germ granules (Figures 3D and 3E), further supporting the notion that the periphery of the organelle represents a site of translation. Ribosomes have also been observed at the periphery of *Drosophila* polar granules shortly after fertilization, but in this case they disappear from these sites prior to the formation of germ cell precursor cells^43–45^. The distinct ribosome localization pattern observed in 10 hpf zebrafish embryos (Figure 3C) was also detected at an earlier stage of germ cell development (Figure S3A), shortly after specification of the germline^32, 46, 47^. Importantly, the peripheral ribosome-rich regions of the granule partially overlap with the peripheral domains in which *nanos3* mRNA clusters reside (Figures 4B and 4C). Interestingly, in addition to the peripheral enrichment of ribosomes, we occasionally detected intra-granule domains that lack Vasa signal. Such domains contained ribosomes and overlapping *nanos3* mRNA (Figure S3B). These observations suggest that the granule-resident Nanos3, a protein essential for germ cell development^25^ (Figure 1A), is translated primarily at the borders of the phase-separated material. To further test this hypothesis we examined an earlier developmental stage, when *nanos3* RNA is translationally silent^46^. In zebrafish embryos, fluorescence of EGFP encoded by maternally supplied *nos* 3’UTR*-*containing mRNA is first detected at 3 hpf^46^, suggesting that translation of *nanos3* mRNA starts at around 2 hpf. We thus determined the distribution of *nanos3* mRNA at the 8-cell stage (1.25 hpf, Figure S3G). At this stage, germ cells have not yet formed, and the germ plasm material appears as a rod-like structure at the cleavage furrows of the blastomeres^32, 47–49^. Using the early germ plasm protein Bucky Ball as a marker^50, 51^, we found that in contrast to later stages when *nanos3* is translationally active, localization of the RNA in 8-cell embryos is restricted to the interior of the germ plasm (Figures S3G and S3H). Whether the germ plasm is already patterned at these stages with respect to the translation machinery remains to be determined.

**Figure 3.**
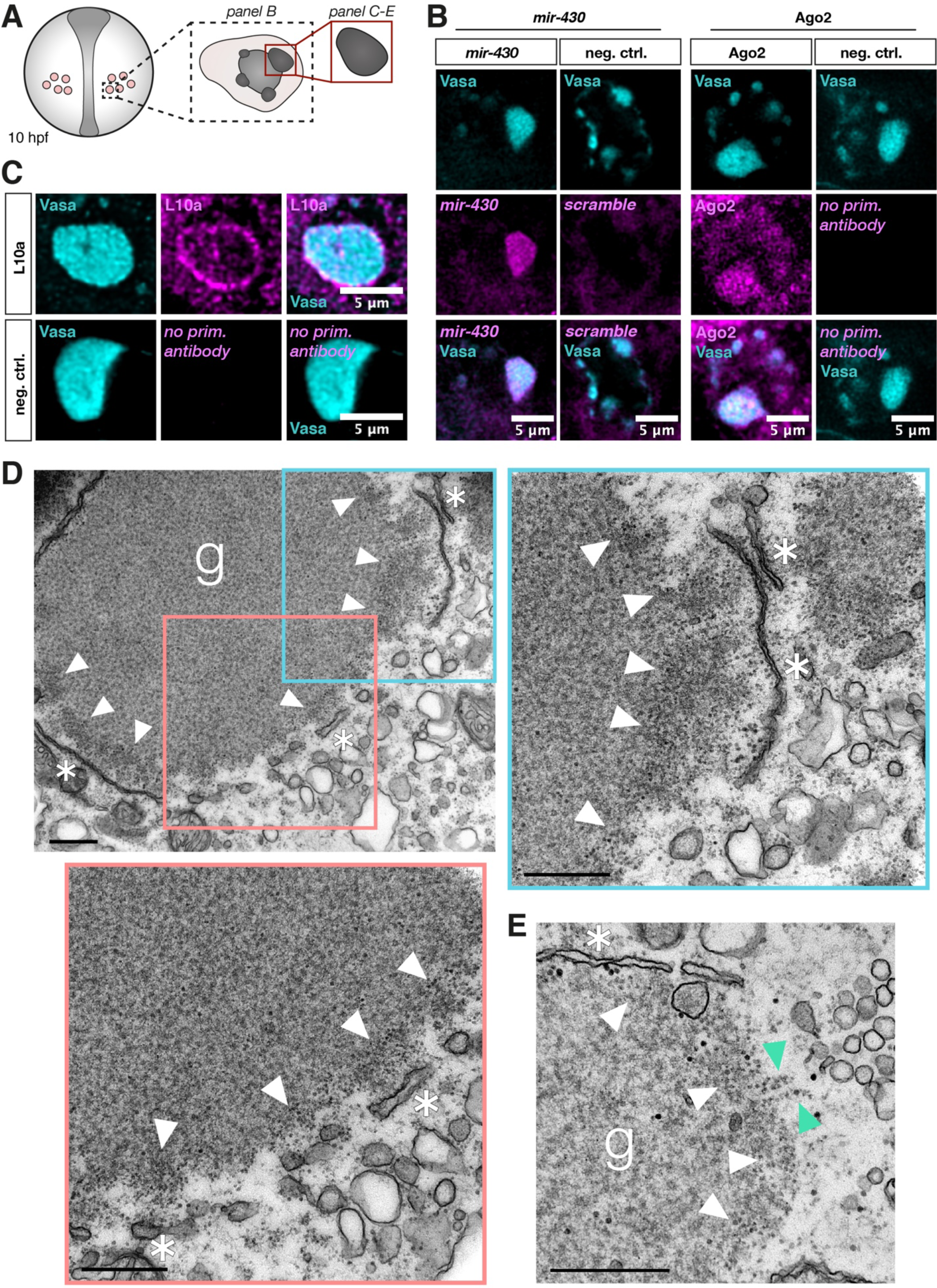
Zebrafish germ granule condensates contain translation-repression factors and exhibit translation activity and enrichment of ribosomes at their periphery. (A) A scheme showing germ cells in a 10 hpf embryo. The black dotted box, and the red box depict the region of interest for the indicated image panels. All panels represent a single confocal plane. (B) Co-visualization of endogenous Vasa protein and endogenous *mir-430* (detected by a miRCURY LNA miRNA probe, negative control = *scrambled* probe) or endogenous Ago2 protein (negative control = no primary antibody) within germ cells. (C) Co-detection of endogenous Vasa protein and the 60S ribosomal protein L10a. Negative control = no primary antibody. (D) Electron microscopy image of a germ granule condensate in a germ cell of a 8-12 hpf embryo. White arrowheads point to enrichment of free ribosomes at the condensate’s periphery, and asterisks mark rough ER membrane structures. Blue and red insets in the upper left image indicate regions shown at a higher magnification in the adjacent images (upper right and bottom left images). g = germ granule. Scale bar = 500 nm. (E) Electron microscopy image of a characteristic structure of polyribosomes (green arrowheads) at the edge of another germ granule. Ribosomes are clearly distinguishable from other cytosolic structures such as glycogen (see Figure S3D). See also Figure S3.

**Figure 4.**
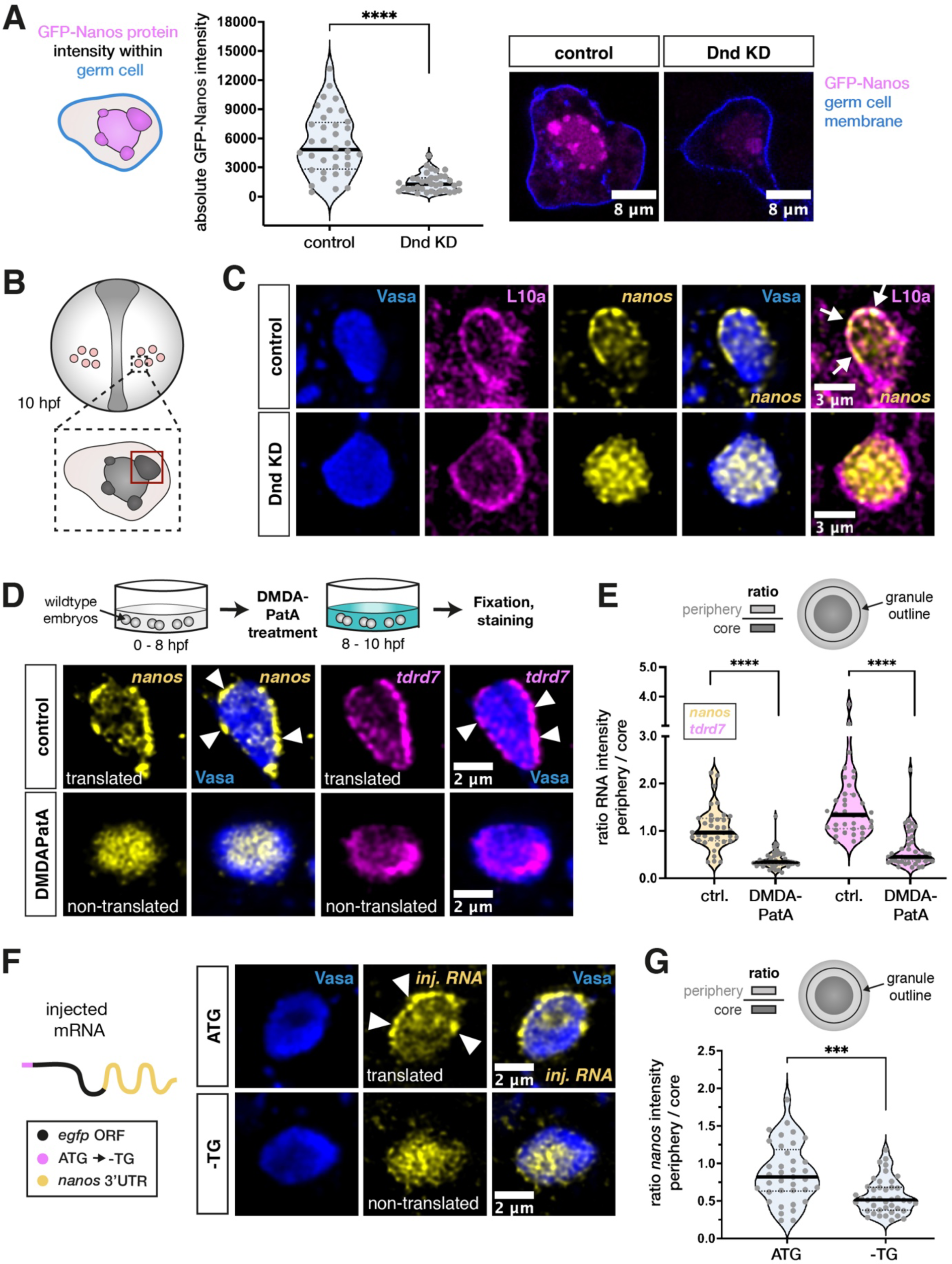
Depletion of Dnd1 or inhibition of translation causes loss of *nanos3* mRNA from peripheral ribosome-rich areas of the germ granule. (A) Evaluation of the mean signal intensity of GFP-Nanos3 protein, encoded by injected *nos* 3’UTR-containing mRNA, within germ cells under control and Dnd KD conditions. The blue cell outline in the scheme represents the border of the area of measurement. Two representative images are shown on the right. Significance was determined using Mann-Whitney *U* test. ****P<0.0001. n (control) = 36, n (Dnd KD) = 36. N = 3. (B) A scheme depicting germ cells in a 10 hpf embryo. The red box outlines the region of interest within the germ cell (large condensate), which was considered for imaging and analysis in panels C to G. All images show one confocal plane of the germ granule. (C) Visualization of endogenous *nanos3* mRNA localization with respect to endogenous Vasa and L10a proteins in germ granules under control or Dnd KD conditions. Arrows point to overlaps of L10a and *nanos3* at the condensate periphery. (D) Localization of endogenous *nanos3* mRNA and *tdrd7* mRNA relative to endogenous Vasa protein in germ granules following control or DMDAPatA treatment. Arrowheads point at clusters of mRNA localized to the granule periphery. (E) Ratio of the mRNA level between the granule periphery and the core of the condensate in control and DMDAPatA-treated cells. Significance was determined using Mann-Whitney *U* test. ****P<0.0001. n (control) = 36, n (DMDAPatA) = 43. N = 3. (F) Localization of injected *nos 3’UTR*-containing mRNA with respect to endogenous Vasa protein in germ granules. The first ATG of the open reading frame (ORF) is either functional (ATG) or mutated (-TG). (G) A graph showing the ratio of the mRNA level between the granule periphery and its core. Significance was evaluated using Mann-Whitney *U* test. ***P = 0.0002. n (ATG) = 52, n (-TG) = 46. N = 3. See also Figure S4.

Together, our findings show that the periphery of germ granules is enriched with ribosomes and ER, and that the translational machinery colocalizes with germline mRNA clusters at the time when the transcripts are actively translated.

### Dnd1 activity is essential for *nanos3* mRNA translation at the periphery of germ granule condensates

To gain further insight into the mechanisms by which Dnd1 controls the function of *nanos3* mRNA within germ granules, we next examined the effect of Dnd1 depletion on the translation of the RNA. Indeed, Dnd1-depleted germ cells show a strong reduction in Nanos3 protein levels, as evaluated by the translation of injected *nos 3’UTR*-containing RNA encoding for a GFP-Nanos3 fusion protein (Figure 4A)^21^. Importantly, the strong reduction in Nanos3 protein synthesis was associated with a loss of the overlap between *nanos3* mRNA and the ribosomes at the germ granule periphery (Figures 4B and 4C). This observation is consistent with the idea that Dnd1 facilitates the association of *nanos3* mRNA with ribosomes and, consequently, to mRNA translation and Nanos3 protein production. Indeed, we found that levels of a EGFP-Tdrd7 fusion protein were not affected by Dnd1 depletion, as expected from the persistence of *tdrd7* mRNA clusters at the granule periphery under these conditions (Figures S4A, 2D and 2E). To determine whether the translational status of the RNA *per se* could be instructive regarding the localization of *nanos3* mRNA, we inhibited translation initiation using a derivative of the natural product pateamine A, des-methyl, des-amino pateamine A (DMDAPatA)^52–55^ (Figure S4B). Intriguingly, under these experimental conditions, we observed a strong depletion not only of *nanos3* mRNA but also of *tdrd7* mRNA from the periphery of the germ granule, with a concomitant increase in the level of the RNAs in the core of the condensate (Figures 4D and 4E). In contrast, Dnd1 protein and ribosomes were mostly maintained at the organelle periphery (Figure S4C). However, the peripheral domains formed by Dnd1-GFP protein appeared broader (Figure S4C, left panels). These findings suggest that a correlation between the translation of RNA molecules and their presence at the border of germ granule condensates is not restricted to *nanos3* mRNA but likely exists for other germline mRNAs as well. In addition to the effect of global inhibition of translation, the specific inhibition of *nanos3* mRNA translation by mutating the first Kozak sequence-containing ATG led to a similar effect. Specifically, *egfp-nos 3’UTR* RNA, in which the translation start codon was mutated, accumulated at the center of the granule and was largely depleted from the periphery (Figures 4F and 4G).

To further test the hypothesis that RNA molecules interact with ribosomes at the granule periphery, we employed the translation elongation inhibitor cycloheximide (CHX), which blocks the elongation of the polypeptide chain and causes stalling of ribosomes on the transcript^56, 57^. Intriguingly, we found that in CHX-treated cells *tdrd7* and *nanos3* RNAs are maintained at the periphery of the granule, with *nanos3* mRNA signal even enhanced (Figures S4D and S4E), suggesting that the interaction of the RNA with the translation machinery helps mainlining it at the edge of the granule.

Together, these findings are consistent with the idea that germ granules are patterned such that mRNA is stored and protected at the granule interior, while translation occurs at the periphery of the condensate, where ribosomes can interact with the transcripts.

### Loss of *nanos3* mRNA from the condensate border in absence of Dnd1 is not a result of transcript degradation, but rather of translocation to the core region

Having shown that the Dnd1 protein is required to facilitate *nanos3* mRNA localization at domains of the germ granule where translation takes place, we proceeded to investigate the potential mechanisms that underly this process. Dnd1 was previously shown to interfere with the function of the micro-RNA (miRNA) *mir-430*, thereby conferring stability to *nanos3* mRNA within the germ cells^22^. Indeed, we found that the total amount of the mRNA is reduced within Dnd1-depleted germ cells at 10 hpf (Figure 5A). Interestingly however, in contrast to the decrease in the overall mRNA levels within the cells, we observed a specific and strong elevation of the amount of *nanos3* mRNA within germ granules in the absence of Dnd1 (Figure 5B). This increase in mRNA abundance suggests that a portion of the accumulated transcripts in the granule core is derived from the cytoplasm. Moreover, our findings indicate that the loss of transcripts from the periphery of the condensate (Figure 2D) is a consequence of translocation to the core domain rather than merely a decay of mRNA at the granule periphery.

**Figure 5.**
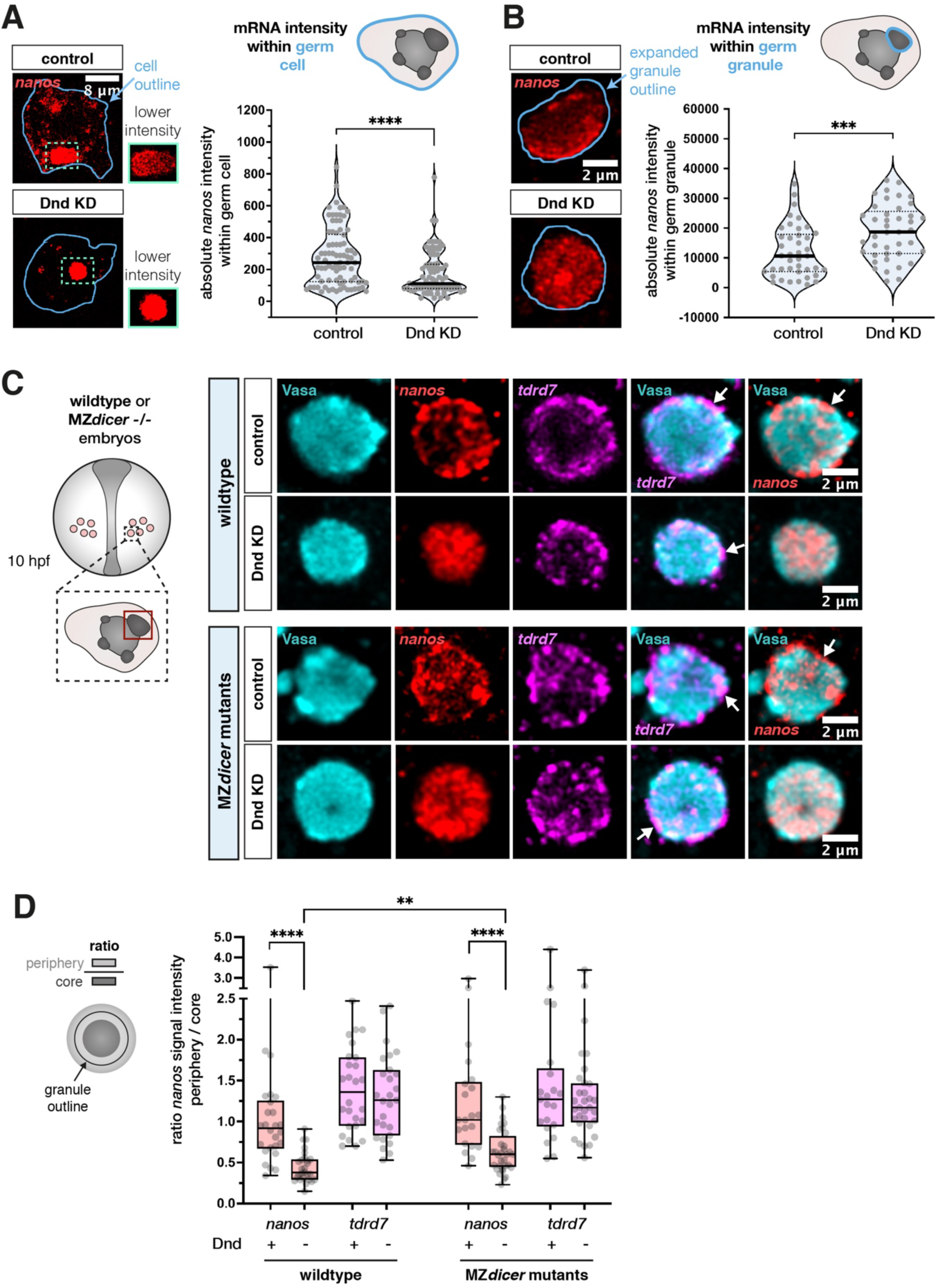
Loss of nanos3 mRNA from the condensate border in absence of Dnd1 is not a result of transcript degradation but rather of translocation to the core region. (A) Analysis of the mean signal intensity level of endogenous *nanos3* mRNA within germ cells under control and Dnd KD conditions. The blue outline in the left panels marks the segmented cell outline, representing the area of measurement that was conducted across a 3D stack. The quantification of the results is presented in the graph. The image panels show representative single-plane confocal images. Green dotted outlines show mRNA accumulation in large condensates of the cells, and the adjacent green boxes show a magnification of the same areas with a lower pixel intensity. Statistical significance was determined using Mann-Whitney *U* test. ****P<0.0001. n (control) = 82, n (Dnd KD) = 86. N = 3. (B) Evaluation of the mean intensity of endogenous *nanos3* mRNA within large germ granules under control and Dnd KD conditions. Left image panels represent one confocal plane of a granule showing mRNA signal. The dilated outline of the segmented condensate (light blue) was used to determine the area of measurement and the graph shows the quantitation of the results. Statistical significance was evaluated using Mann-Whitney *U* test. ***P = 0.0007. n (control) = 41, n (Dnd KD) = 41. N = 3. (C) Localization of endogenous *nanos3* and *tdrd7* mRNA in wildtype (upper two rows) and MZ*dicer-/-* embryos (lower two rows) under control and Dnd KD conditions, with a single confocal plane presented. Arrows point at mRNA enrichment at the condensate periphery. The red box in the left scheme outlines a region of interest within the germ cell (large condensate), which was considered for imaging and quantitative analysis presented in (D). (D) The ratio between the mRNA level in the granule periphery and its core for endogenous *tdrd7* and *nanos3* mRNAs in wildtype and MZ*dicer* embryos. Significance was calculated using Mann-Whitney *U* test. **P = 0.0021, ****P< 0.0001. Wildtype n (control) = 26, n (Dnd KD) = 27. MZ*dicer* n (control) = 21, n (Dnd KD) = 32. N = 3.

Consistent with the idea that loss of *nanos3* RNA from the granule periphery at these stages results from relocation of the transcript, rather than from degradation, we analyzed the effect of Dnd depletion in maternal zygotic (MZ) *dicer* mutants, which lack mature miRNAs^58, 59^ (Figures 5C and 5D). Indeed, the depletion of *nanos3* mRNA from the condensate periphery upon Dnd KD was observed also in these mutants, showing that it occurs independently of miRNA activity. Rather, our findings suggest that the positioning of *nanos3* mRNA within the condensates is primarily controlled by its interaction with the active translation machinery. Together, we conclude that inhibition of Dnd1 renders *nanos3* mRNA inaccessible to the translation machinery, leading to its accumulation in the granule core, and consequently to a reduction in Nanos3 protein levels.

## DISCUSSION

Our findings reveal how phase-separated vertebrate germ granules are organized concerning the spatial distribution of RNAs, proteins and the translation machinery. Importantly, we found that following the formation and segregation of germ plasm into the future germ cells^6, 60^, Dnd1 plays a key role in controlling the accessibility of granule-localized RNA to the translation machinery, and we present the relevance of this arrangement for cell fate and germline development (Figure 6).

**Figure 6.**
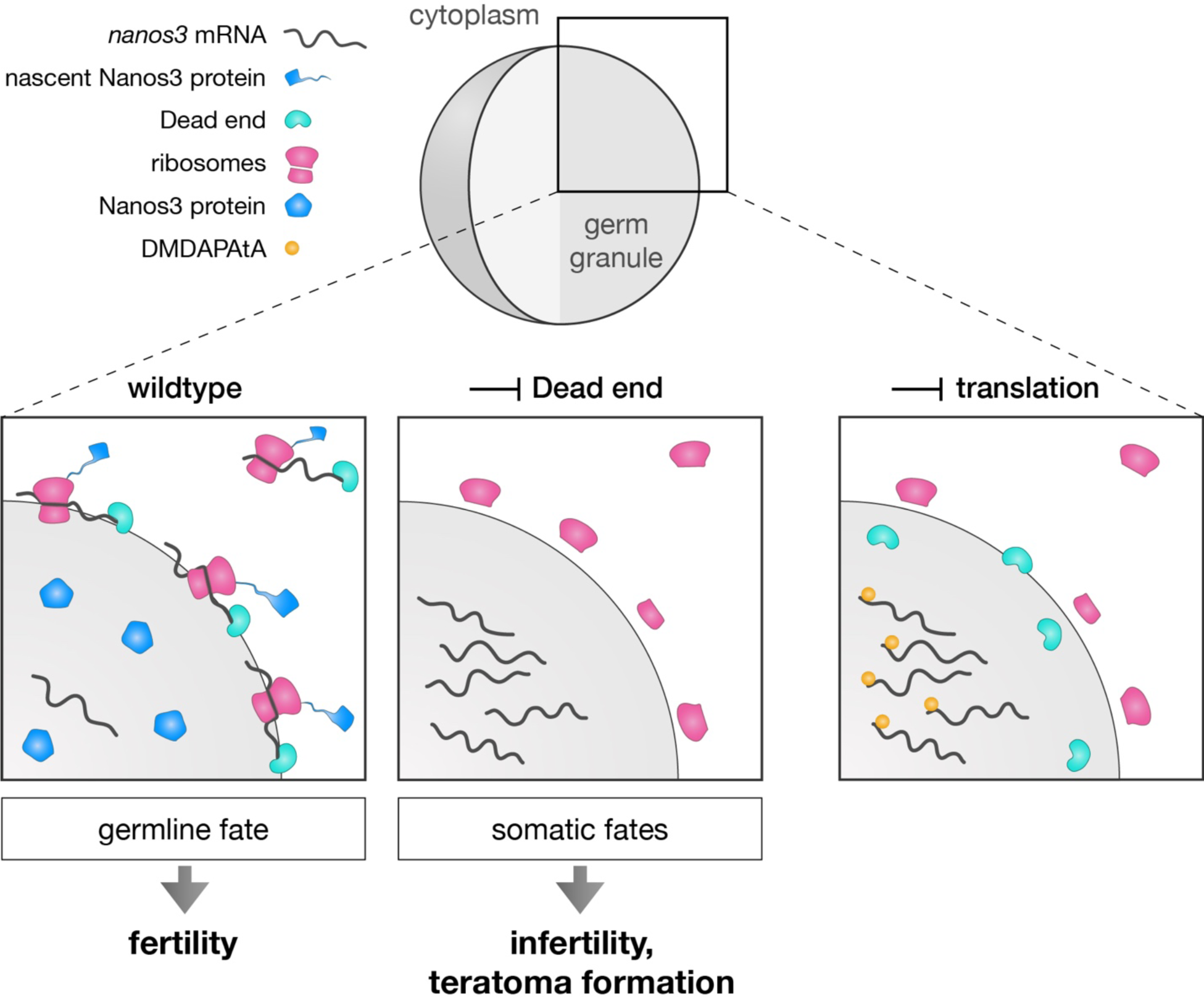
Model for regulation of *nanos3* localization and function by Dnd1 in germ granules. In wildtype cells (left panel), *nanos3* mRNA associates with Dnd1 at the periphery of the germ granule, where ribosomes reside. At this site, an active translation complex forms on the mRNA, resulting in Nanos3 protein synthesis. Under these conditions, the germ cells maintain their fate, and a fertile organism develops. Upon depletion of Dnd1, *nanos3* mRNA translocates to the core of the condensate, where it is inaccessible to the ribosomes, resulting in lack of Nanos3 protein synthesis (middle panel). Under these conditions, the germ cells acquire somatic fates, leading to infertility. Upon inhibition of translation initiation complex assembly by DMDAPatA treatment (right panel), *nanos3* mRNA is not bound by ribosomes and translocates to the granule core.

In many organisms, post-transcriptional regulation and properties of phase-separated condensates are key for specifying and preserving the fate of germ cells^61, 62^. During early germ cell development, RNA translation must occur rapidly to ensure the segregation of the germline from the soma, but at the same time a pool of RNAs should be maintained for translation at later stages. Accordingly, recent studies show that translationally repressed mRNAs accumulate within germ granules, where they are stored and protected from degradation^10, 11^. The initiation of translation of such RNA molecules would require interactions between the transcripts and the translation machinery outside of the organelles. Our study reveals the presence of ribosomes and ER at the periphery of phase-separated germ granules, and as we show for *nanos3* mRNA and *tdrd7* mRNA, these germline mRNAs appear to interact with ribosomes at the condensate border. The presence of the transcripts at this region is correlated with their translation, and with maintenance of the germline fate (Figure 6). At the same time, our results are consistent with the idea that RNA located at internal domains of the granule is stored and undergoes translation at later stages of germ cell development.

As exemplified here for *nanos3* RNA, our findings suggest that germ granules are bifunctional organelles. We suggest that a fraction of the mRNA is localized to and translated at the organelle’s periphery, while the rest is preserved and protected from degradation at the core of the granule, for use at later stages. Thus, we consider the core-periphery patterning of germ granules as a mean for controlling the level and time of mRNA translation in the cells, thereby ensuring proper germline development at different stages. The requirement for such control mechanisms could be especially relevant in species such as zebrafish, *Xenopus*, *Drosophila* and *C. elegans*, where germ cells are specified and maintain their fate for extended time using maternally provided RNA and protein molecules^12, 63, 64^. Relevant for this point, in the *C. elegans* model it was found that P granules are largely dispensable for fertility of the worm^10^. These observations could reflect differences among species, and the extent to which maternal transcripts have to be maintained and used. Investigating these points would require a detailed analysis of the fate and dynamics of RNA molecules within the condensate and of the amount of protein translated at the granule surface. In zebrafish, the generation and maintenance of a sub-granule organization of molecules and structures (e.g., ribosomes) occurs in relatively large condensates (up to 6 µm in diameter as compared with 0.5 µm in *Drosophila* at early germline development^65^). It would thus be interesting to determine to what extent the mechanisms presented here play similar roles in other organisms.

Importantly, the data we present points at a role for RNA-binding proteins in RNA localization within the granule and control over translation. In the case of *nanos3* mRNA, we show that in the absence of Dnd1 function, reduced *nanos3* RNA translation leads to loss of germ cell fate. While the precise mechanisms by which Dnd1 controls *nanos3* localization are unknown, our findings suggest that the formation of an active translation complex is required for maintaining mRNA clusters at the condensate periphery. Recent findings in *C. elegans* link P granule localization of *nanos* mRNA to translation repression^10, 11^. Similarly, association of repressors of *nanos3* translation in Dnd1-depleted germ cells could maintain the transcript at the core of the granule.

Consistently, in *Xenopus*, Dnd1 was shown to prevent interaction of translational repressor eIF3f with *nanos* mRNA^66^. It would be interesting to determine whether a similar mechanism functions in zebrafish as well. In addition, future studies should be directed towards determining the role of proteins other than Dnd1 in controlling the localization and function of other RNA molecules such as that encoding for Tdrd7.

Our study reveals RNA translation activity at the periphery of phase-separated germ granules (Figure 6). Nevertheless, we did not determine the contribution of translation at the granule surface relative to that in the cytoplasm at different time points. Indeed, a recent study in *C. elegans* found that in absence of germ granules, translational activation of germline transcripts still occurs^10^. Future studies should thus focus on determining the importance of each translation site for regulating the protein levels required for proper germline development.

### Limitations of the study

Of note, our study here has certain limitations. We provide information regarding the control over *nanos3* RNA function in germ granules and the role of the Dead end protein in this context. As exemplified by *tdrd7* RNA, other RNAs are regulated differently. Identifying molecules that control the activity and localization of other RNAs that reside within germ cell granules will provide a more comprehensive understanding of the mechanisms controlling the function of the condensates. As mentioned above, we did not determine the proportion of translation occurring at the periphery of the granules relative to that at other parts of the cell. Evaluating this parameter in germ cells at different phases of their development will help assigning roles for this level of regulation in processes culminating in gamete formation. Last, it would be interesting to determine to what extent the mechanisms we identify here are relevant for other species in which early germ cell development relies on germ plasm components.

## Supporting information

Westerich_et_al_Supplemental

## ACKNOWLEDGMENTS

We thank Celeste Brennecka for editing and Michal Reichman-Fried for critical comments on the manuscript. We thank Ursula Jordan, Esther Messerschmidt and Ines Sandbote for technical assistance. This work was supported by funding from the University of Münster (KJW, KT, ER, AG, TGT, JS, MG), the Max-Planck for Molecular Biomedicine (DZ), the German Research Foundation grant CRU 326 (P2) RA863/12-2 (ER), Baylor University (KH, DR) and the National Institutes of Health grant R35 GM 134910 (KH, DR). We thank the referees for insightful comments that helped improve the manuscript.

## AUTHOR CONTRIBUTIONS

E.R. supervised the project. K.J.-W. and E.R. designed the work and wrote the manuscript. K.J.- W. performed all of the experiments except for the following: T.G.-T. quantified the cell shape of Nanos-depleted germ cells and A.G. acquired the respective microscopy images; J.S. analyzed the distribution of domains with enriched molecules in germ granules; K.T. generated MZ*dicer* mutants; M.G. prepared the samples for the electron microscope and D.Z. performed the electron microscopy studies; M.Z. synthesized DMDAPatA, K.H. and D.R. supervised the synthesis of the translation initiation inhibitor, des-methyl des-amino pateamine A (DMDAPatA). All authors read and approved the final manuscript with minor revisions.

## STAR METHODS

### Key Resources Table

Individually provided.

### RESOURCE AVAILABILITY

#### Lead contact

Any information or requests for resources and reagents should be directed to the Lead Contact, Erez Raz.

#### Materials availability

All plasmids generated in this study are available from the lead contact.

#### Data and code availability

- No large-scale datasets have been generated in this study. Microscopy data reported in this paper will be shared by the lead contact upon request. Accession numbers are listed in the key resources table.
- This paper does not report original code.
- Any additional information required to reanalyze the data reported in this paper is available from the lead contact upon request.

### EXPERIMENTAL MODEL AND STUDY PARTICIPANT DETAILS

#### Zebrafish strains and handling

The following lines were used to obtain zebrafish (*Danio rerio*) embryos: wild type of the AB or AB/TL genetic background, *cxcr4b^t^*^2603567^, *medusa^NYO^*^45^ ^68^, transgenic *kop-egfp-f-nos-3′UTR*^69^ and transgenic *buc:buc-egfp*^51^ in a *buc^p1^*^06r*e*^ ^50^ mutant background. Maternal zygotic (MZ) *dicer* embryos (*dicer^hu715^*^70^) were generated by conducting a germline replacement as described previously^58^. Embryos were raised in 0.3× Danieau’s solution (17.4 mM NaCl, 0.21 mM KCl, 0.12 mM MgSO_4_·7H_2_O, 0.18 mM Ca(NO_3_)_2_) at 25°C, 28°C or 31°C. The fish were handled according to the regulations of the state of North Rhine-Westphalia, supervised by the veterinarian office of the city of Muenster.

### METHOD DETAILS

#### Cloning of DNA constructs

To follow mRNA using the PP7 detection system in live embryos, the sequence of 24 PP7 stem loops^71^ was cloned into a Nanos3-nos 3’UTR plasmid^25^ downstream of the open reading frame (ORF) sequence. The Dnd-mScarlet-nos 3’UTR plasmid was generated by replacing the GFP sequence of a Dnd-GFP-nos 3’UTR plasmid^72^ with that of mScarlet. The *nos 3’UTR*-containing mRNAs used in Figures 2F, 2G, 4F, 4G, S4B and S4C all carry an EGFP sequence in the ORF to allow for detection by RNAscope probes. For the generation of a non-translatable *nos 3’UTR*-containing mRNA used in Figures 4 F and 4G, a 1-base pair deletion was introduced into the KOZAK-flanked ATG start codon of an EGFP coding sequence located upstream of the *nos 3’UTR* (EGFP^-TG^-nos 3’UTR). For this purpose, site-specific mutagenesis was employed^73^. For the generation of a translatable *nos 3’UTR*-containing mRNA used in Figures 2F, 2G, 4F and 4G, the sequence encoding for amino acids 2-15 was deleted from a full-length EGFP sequence-containing plasmid employing a Restriction free cloning approach^74^ to prevent fluorescence emission of the EGFP protein (EGFP^Δ2–15^-nos 3’UTR). This step was required to simultaneously visualize multiple RNA transcripts without emission of the EGFP protein. To generate a *nos 3’UTR*-containing mRNA lacking all start codons in the ORF used in Figures S4B and S4C, all ATGs within the EGFP^Δ2–15^-nos 3’UTR sequence were mutated to GTGs. Guanine was chosen to replace Adenine based on the structural similarity of the two bases. The respective ORF sequence carrying the ATG > GTG conversions was synthesized by Eurofins Genomics and cloned into a *nos 3’UTR*-containing plasmid. To generate the GFP-globin 3’UTR plasmid for global GFP expression in zebrafish, the GFP sequence was cloned upstream of a *Xenopus globin 3’UTR*.

#### Microinjection into zebrafish embryos

Capped sense mRNAs were synthesized using the mMessage mMachine kit (Ambion) according to the protocol of the manufacturer. 1 nL of mRNA and / or Morpholino-containing solution were injected into the yolk of 1-cell stage embryos. To mark the membrane of somatic cells, 45 ng/µl of *mcherry-f’-glob 3’UTR (A709)* were injected^75^. For visualization of protein within germ granules, *vasa-gfp-vasa 3’UTR* mRNA *(291)*^76^ was injected at a concentration of 100 ng/µl, *dnd-gfp-nos 3’UTR (516)* mRNA^72^ at 180 ng/µl, *mgfp-nanos-nos 3’UTR (356)* mRNA at 90 ng/µl^25^ and *egfp-tdrd7-tdrd7 3’UTR* at 150 ng/µl^13^. For the evaluation of Dnd protein and *nanos3* mRNA movement, 140 ng of *nanos-nos 24xPP7 3’UTR (D016)* and 8 ng of *nls-ha-tdpcp-yfp-glob 3’UTR (C987)*^77^ were co-injected with 100 ng of *dnd-mscarlet-nos 3’UTR (E513)*. To visualize *nos 3’UTR*-containing mRNA localization within germ granules, *egfp^-TG^-nos 3’UTR, egfp^Δ2-15-nos 3’UTR^* and *egfp^Δ2-15_mut-ATG^-nos 3’UTR (D366, E801, E845)* transcripts were injected at an equimolar concentration of 100 ng. For the evaluation of DMDAPatA drug efficiency, *gfp-globin 3’UTR* mRNA (B289) was injected at a concentration of 50 ng/µl. Nanos3 Morpholino (TGAATTGGAGAAGAGAAAAAGCCAT) was injected at a concentration of 60 µM^25^, Dnd Morpholino (GCTGGGCATCCATGTCTCCGACCAT) at 20 µM or 30 µM (Figure 4A)^72^ and Cxcl12a MO (TTGAGATCCATGTTTGCAGTGTGAA) at 200 µM^29^.

#### RNA detection and immunostaining of whole-mount embryos

For all treatments, embryos were fixed in 4% PFA overnight at 4°C (6 hpf and 10 hpf stage) or for 2 h at room temperature (RT, 10 hpf stage) under slow agitation. Embryos were subsequently rinsed in 1x PBT, dechorionated and transferred to and stored in 100% methanol. Detection of *mir-430* RNA in whole-mount embryos was performed using an in-situ hybridization procedure optimized for miRNA detection as previously described^78, 79^. Dual labeled miRCURY LNA miRNA detection probes of *dre-mir-430a-3p* and *scramble-miR* (Qiagen, catalogue no. YD00613610-BCO and YD00699004-BCO) were hybridized at a concentration of 62.5 nM overnight at 45°C. To co-visualize proteins with mRNAs within germ cells, an RNAscope *in situ* hybridization procedure was conducted using the RNAscope Multiplex Fluorescent Reagent Kit (ACD Bio) as previously described^80^. RNAscope probes Probe Diluent (ACD Bio 300041), Dr-nanos-C2 (ACD Bio 404521-C2), Dr-tdrd7-C2 (ACD Bio 536551-C2), EGFP-C1 (ACD Bio 400281-C1) and Dr-nanos3-CDS-C3 (ACD Bio 431191-C3) were hybridized overnight at 40°C. The embryos were then subjected to an immunostaining procedure as previously described, with a few modifications^80^. Briefly, embryos were permeabilized in 1x PBS containing 0.1% Tween-20 and 0.3% Triton X 100 (Carl Roth) for 1 h at RT and subsequently incubated in blocking solution (0.3% Triton X 100 and 4% BSA in 1x PBS) for 6 h at RT. Incubation with primary antibodies was then performed for 24 h at 4°C and incubation with secondary antibodies was done overnight at 4°C. The treatment with a GFP-targeting antibody was performed to enhance the fluorescent signal of expressed GFP-tagged Dnd protein. For the detection of Ago2 protein in germ cells, embryos were incubated in Dents solution (20% DMSO in methanol) for two days at 4°C, rehydrated in increasing amounts of 1x PBT in methanol (25%, 50%, 75% and 100%), re-fixed in 4% PFA for 20 minutes at RT and rinsed in 1x PBT. Embryos were then incubated in 1x PBT + 0.3% Triton X 100 for 1 h at RT and subsequently incubated for 5 h in blocking solution (1 x PBT + 20% goat serum + 5% DMSO) at RT. Subsequent treatment with primary and secondary antibodies was conducted as above.

Primary antibodies used: anti-Vasa (rabbit polyclonal, 1:400 dilution, GeneTex catalogue no. GTX128306), anti-GFP-1020 (polyclonal chicken, 1:600 dilution, Aves labs: SKU GFP-1020), anti-RPS2 (polyclonal rabbit, 1:300 dilution, GeneTex catalogue no. GTX114734), anti-RPS6 (polyclonal rabbit, 1:100 dilution, GeneTex catalogue no. GTX113542), anti-eIF4G (monoclonal mouse, 1:250 dilution, Sigma catalogue no. SAB1403762-100U), anti-PABP (polyclonal rabbit, 1:250 dilution, ThermoFisher catalogue no. PA5-17599), anti-eIF4E (polyclonal rabbit, 1:250 dilution, ThermoFisher catalogue no. PA5-86047), anti-RPL10a (monoclonal mouse, 1:100 dilution, Sigma catalogue no. WH0004736M1-100UG), anti-Ago2 (monoclonal mouse, 1:500 dilution, Abcam catalogue no. ab57113). Antibodies were selected based on reported activity in zebrafish (anti-Vasa, anti-RPS2, anti-RPS6, anti-PABP, anti-Ago2^79^) or high sequence conservation of the target sequence among the target species and zebrafish (anti-eIF4G, anti-eIF4E and anti-RPL10a).

Secondary antibodies used: to visualize GFP, a 1:600 dilution of a 488 nm anti-chicken secondary antibody was employed (Jackson Immuno-Research); in all other cases, secondary antibodies from Thermo Fisher Scientific, catalogue no. A-11031, A-11034, A-11036 were employed using a 1:500 dilution for visualizing Vasa and a 1:1000 dilution for visualizing the remaining proteins.

#### Electron microscopy (EM) of germ granules in zebrafish germ cells

To visualize ribosomes within germ cells of the zebrafish embryo at 8-12 hpf, laser-mark assisted Correlative Transmission Electron Microscopy (TEM) was applied, using GFP-guided identification of germ cells within the embryo tissue as previously described^81^. 60 nm thick ultrathin sections were analyzed and the images presented show representative structured germ granules. Four independent experiments were conducted and all imaged cells exhibited similar features with respect to their ultrastructural organization. Germ cells could be clearly identified by the presence of germ granule condensates. Subcellular structures including nucleic acids were visualized by a combination of uranyl acetate and lead staining. Lead staining alone was employed to highlight glycogen structures within the cell, making them better distinguishable from ribosomes using serial, consecutive sections.

#### Microscopy and image processing in zebrafish embryos

Images of germ granules in germ cells were acquired using a confocal microscope (LSM 710, Zeiss) equipped with 405 nm, 488 nm, 561 nm and 633 nm lasers and a 40x W N-Achroplan and a 63x W Plan-Apochromat objective (Zeiss). The ZEN software (Zeiss, version 2010B SP1, 6.0) was used to control the microscopy setup. Acquisition of different samples was conducted using the same pinhole size, optical slice thickness, gain and offset parameters. For images showing protein and RNA localization within germ granules, images were acquired with varying laser intensities (except for data presented in Figures 4A, S4A, 5A and 5B) due to variability in germ cell depth within the organism and in germ granule size. This facilitated an optimal resolution without over exposure, which is required for accurately resolving mRNA structures. In this case, the mRNA signal intensities across the germ granule were expressed in relative amounts to compare the mRNA distribution among samples. For the 3D granule layer analysis in Figures 1E, 2E and S2C, images of granules were acquired with a z-distance of 300 nm. To measure signal intensities of *nanos3* mRNA within germ cells (Figure 5A), images were acquired with a z-distance of 1 µm. For the time lapse data shown in Figure 2C and Movie S1, stacks with a z-distance of 500 nm were acquired with a Yokogawa CSU-X1 Spinning Disk microscope equipped with a Piezo-controlled 63x W Plan-Apochromat objective (Zeiss) and connected to a Hamamatsu Orca Flash 4.0 V3 camera. The VisiView software (Visitron, version 4.0.0.14) was used to control the setup of the microscope. For the verification of Pateamine A drug efficiency, data was acquired with a 5x EC Plan-Neofluar objective (Zeiss) on the Spinning Disk. EM images of germ granules were acquired with a Tecnai 12 transmission electron microscope (Thermofisher, Eindhoven, NL) equipped with a biotwin lense and connected to a veleta CCD camera (EMSIS, Münster, Germany). Images acquired with the confocal or Spinning Disk microscope were deconvolved using the Huygens Professional version 19.04 software (Scientific Volume Imaging, The Netherlands, http://svi.nl), employing the CMLE algorithm, with a maximum of 40 iterations.

#### DMDAPatA and CHX treatment of zebrafish embryos

To block translation of mRNAs, 8 hpf zebrafish embryos were incubated for 2 h at 28°C in Danieau’s medium containing either 10 µM DMDAPatA, a derivative of Pateamine A (synthesized by the lab of Daniel Romo, Baylor University), 50 µM CHX (Sigma) or DMSO. Embryos were then fixed in 4% PFA overnight at 4°C under slow agitation. Embryos were subsequently rinsed in 1x PBT, dechorionated and transferred to and stored in 100% methanol until subjected to RNAscope and immunostaining procedures. For the evaluation of DMDAPatA drug efficiency, embryos were injected with 50 ng/µl of *gfp-globin 3’UTR* mRNA at the 1-cell stage and incubated with DMDAPatA or DMSO from the 16-cell stage until 5 hpf. Global expression of GFP was then compared between drug- and DMSO-treated embryos. Since DMDAPatA treatment stops embryonic development, the incubation period of these embryos was timed by monitoring the development of DMSO-treated control embryos, as described previously^82^.

### QUANTIFICATION AND STATISTICAL ANALYSIS

#### Evaluation of somatic cell shape acquisition by germ cells

To analyze the effect of Nanos3 depletion on the trans-fating potential of germ cells, *medusa^NYO^*^45^ embryos that lack expression of the guidance cue Cxcl12a were employed. In such embryos germ cells are localized at ectopic locations. Control or Nos Morpholino were injected at the 1-cell stage into these embryos, and the shape of germ cells was assessed at 24 hpf based on EGFP-F’ expression. To have a clear definition of a somatic cell shape, germ cells were compared to three common somatic cell types (muscle, notochord or neuronal cells) and counted as acquiring a somatic cell shape only if their outline matched with either of these cell types. The actual proportion of Nos KD germ cells that are transfated is therefore likely to be higher than reflected in the graph presented in Figure 1B. Cell proportions were calculated by measuring the ratio between the number of germ cells with muscle, notochord or neuronal cell shape and the number of all germ cells. For the images shown in Figure 1A, a Morpholino antisense oligonucleotide was employed to inhibit translation of the Cxcl12a-encoding mRNA.

#### Analysis of the distribution of clusters of enriched protein or RNA material within germ granules

The analysis of the distribution of clusters of enriched protein or RNA material within germ granules presented in Figures 1G and 2F was conducted on cross-sections of the condensates, which were divided in core and peripheral regions. The peripheral region also includes the space directly surrounding the germ granule. To define the regions, the following analysis was performed using the Fiji software (National Institutes of Health, version 2.0.0-rc-43/1.51a): A binary image of the Vasa granule marker channel was created using an auto intensity threshold, followed by a duplication of the image. One of the duplicated versions was dilated (3 iterations) to upscale the granule area, the other version was eroded (5 iterations) to downscale it. A region of interest (ROI) was defined for the dilated granule area and inversed. This inversed ROI was combined with the ROI of the eroded granule area. The combined ROI was inversed again, resulting in a layer-like space covering the periphery of the granule. Another ROI was created around the eroded signal to cover the core region.

The following analysis was then performed in ImageJ (Version 1.53t), to determine the distribution of clusters of enriched protein or RNA material within the condensate core and periphery: Image processing was conducted for generating masks of bright clusters within the image area. 0.05% of pixels were saturated, the signal histogram was normalized over the 16-bit signal range, and a median filter was applied at a pixel radius of 2.5. The processed images were converted to a mask by applying a threshold of 5% of the highest intensity counts on the histogram. From the resulting mask, subsequent interior and periphery-specific masks of bright clusters were generated by assigning the corresponding ROIs. These masks were applied to the original image to measure the dimensions and intensity of bright clusters in the interior and periphery of granules. Finally, the total area of clusters within the granule core or periphery was calculated and normalized to the area of the respective region. The percentage of area covered with clusters in the core and periphery was then calculated and presented in the graph.

#### 3D layer analysis and volume measurement of germ granules

The analysis of mRNA and protein distribution within defined regions of the germ granule was conducted by employing a 3D image segmentation and processing workflow using the Software Imaris (Imaris 8.4.2, Oxford Instruments). The following steps were carried out to divide the germ granule volume into 3D layers (see also Fig. S1B): Germ granules imaged in the form of confocal stacks with a z-distance of 300 nm were pre-selected based on a minimum radius length of 750 nm and a clearly visible mRNA accumulation. A 3D segmentation of the granule was conducted based on an automated intensity thresholding of the Vasa protein signal. Subsequently, the minor axis length of the germ granule was determined. The ‘Increase Surfaces’ plugin (Imaris XTension, Dominic Filion) was then used to expand or shrink the segmented 3D granule surface by the same increment (a quarter of the minor axis length). The plugin thus generates different-sized surfaces, while preserving the shape of the granule. The RNA or protein signal within each of these surfaces was extracted, and a subsequent subtraction of the extracted signals resulted in the division of the molecules into 3D granule layers. Importantly, the outermost layer covers the space directly surrounding the granule, thus considering molecules that are associated with the outer granule surface. After evaluating the integrity of the layers, the sum of the molecule signal intensity was extracted from each layer and normalized to the volume of the respective layer. The share of signal relative to the total amount in the granule was then calculated and presented in percentage. To evaluate germ granule size, the volume computed from the segmented Vasa protein signal was extracted. For the comparison of mRNA localization among different germ granule sizes, the ratio of normalized mRNA intensity in the peripheral layer to that in the core was determined. The dataset was divided into three subgroups according to the radius of the germ granules.

#### 3D co-localization analysis among germ granule components

For the co-localization analysis of different germ granule components, the Coloc module of the software Imaris was used. To ensure an unbiased and reliable analysis, the following steps were implemented: a surface was created based on the signal of one of the two granule components (e.g., *nanos3* mRNA) and the automatically computed surface intensity threshold value was extracted. The same process was repeated for the second granule component (e.g., *tdrd7* mRNA). The two extracted values were then multiplied by a factor of 1.7 (determined in pre-trials) and served as a minimum intensity threshold for co-localization. Values extracted from the co-localization analysis were the Pearson’s Coefficient (PC) (graphs in Figure S2B) and the amount of material co-localized above threshold (region of overlap in the graph in Figure S1C).

#### Measurement of the periphery-core ratio of mRNA within germ granules

To quantify the relative distribution of mRNA in germ granules as shown in Figures 2G, 4E, 4G, 5D, S3H and S4E, a ratio of the relative mRNA abundance in the granule periphery and core region was determined. The regions were generated using the Fiji software as described above (Method section ‘Analysis of the distribution of clusters of enriched protein or RNA material within germ granules’). Subsequently, the ratio of the mean mRNA intensity in the segmented granule periphery region to that in the segmented core region was determined.

#### Evaluation of absolute RNA and protein signal intensities within germ cells

RNA and protein signal intensity measurements shown in Figures 4A, S4A, 5A, 5B and S4D were conducted with the Fiji software. For the measurement of mRNA intensities within germ granules (Figure 5B), a cross section of a granule was segmented based on the Vasa protein signal as described above. The segmented granule outline was dilated with 3 iterations to upscale the granule area and thereby incorporate all peripheral mRNA signal. The dilated outline was used as a ROI to measure the mean intensity of the mRNA in this region. To measure mRNA intensities in the whole germ cell volume (Figure 5A), cells were segmented across a 3D image stack of 1 µm-distanced slices based on a membrane marker signal that had been pre-processed with a median filter (2 x 10) for noise reduction. The resulting outlines were used as a ROI to extract the mean intensity of mRNA signal in the germ cell. Nanos3 and Tdrd7 protein levels within germ cells (Figures 4A and S4A) were measured employing the same approach, but focusing on a single cross section of the cell that covered the area of nucleus and germ granules.

#### Statistical analysis

The Prism software (version 9.4.0, Graphpad) was used to perform statistical analysis. Three biological replicates were conducted for each experiment, and the data pooled into one sample. Significance was evaluated using the Mann-Whitney *U* test for non-parametric data sets. For comparing RNA localization in different-sized granules, a Kruskal-Wallis test was applied to determine significance among three groups. Embryos and cells were analyzed with non-biased approaches.

## LEGENDS TABLES AND MULTIMEDIA FILES

**Movie S1. Example of the spatial organization of molecules within zebrafish germ granules, related to Figure 1**

The movie shows a 3D germ granule as visualized by Vasa protein signal. The Vasa surface is segmented to outline the shape of the granule, and to visualize the localization of germline mRNAs relative to it.

**Movie S2. *nanos3* mRNA and Dnd1 protein clusters are motile but remain in proximity, related to Figure 2**

Maximum intensity projection showing the mobility of protein and mRNA clusters within a large germ granule. An injected *nos 3’UTR*-containing mRNA carrying PP7 stem loops was detected by the NLS-PCP-YFP coat protein (cyan). Dnd-mScarlet protein is shown in magenta. Arrowheads point to mRNA and protein clusters within the granule that remain in proximity over time. Time interval = 1.5 seconds. The maximum intensity projections are comprised of 4 focal planes with a z-distance of 500 nm.

**Table S1. Oligonucleotides used in this work, related to STAR Methods**

