## Supplementary material for "Spatial organization and function of RNA molecules within phase-separated condensates are controlled by Dnd1": Westerich_et_al_Supplemental

### SUPPLEMENTARY MATERIALS

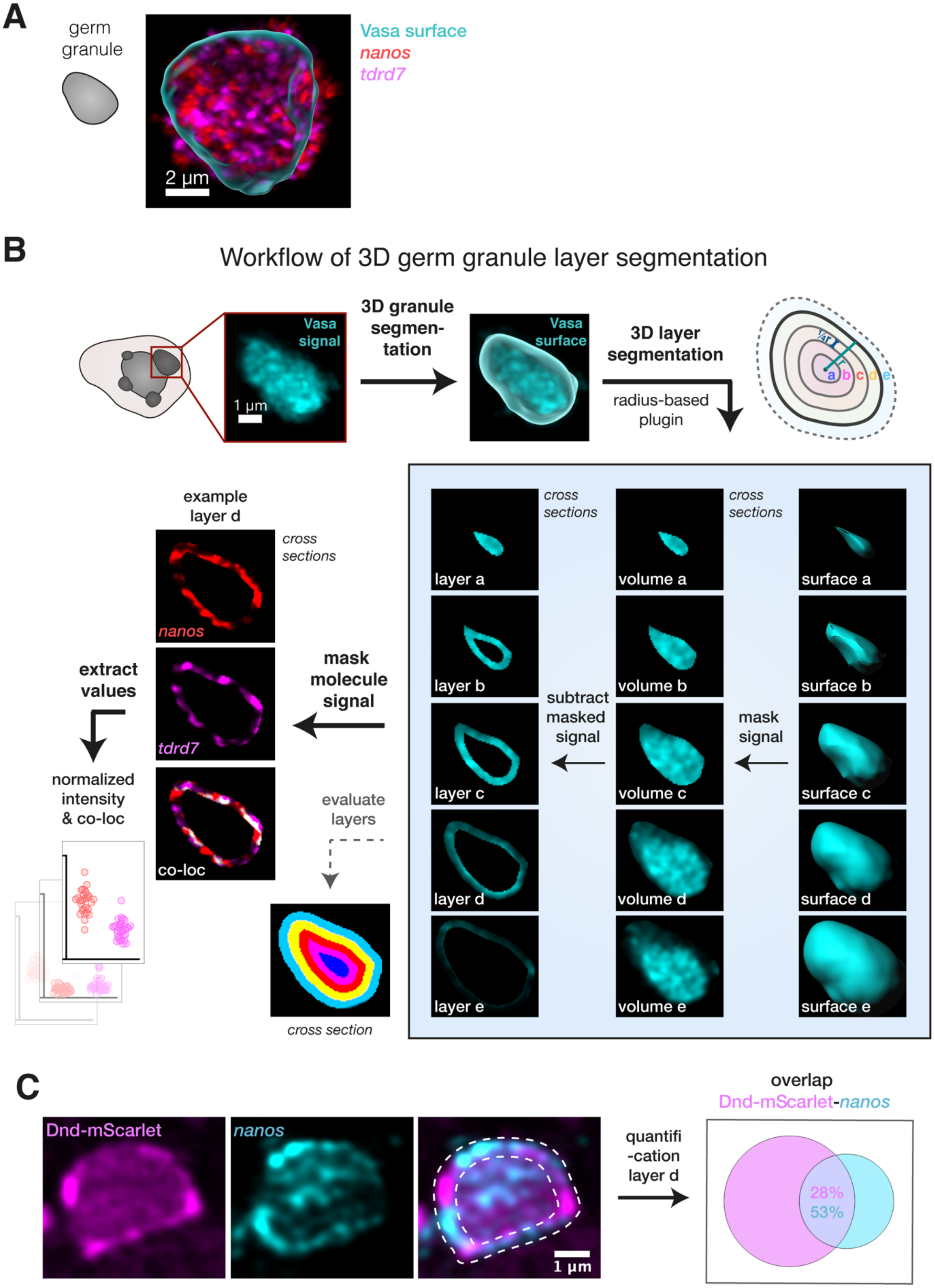

**Figure S1. 3D germ granule layer analysis in zebrafish embryos, related to Figure 1.**
(A) Example of *nanos3* and *tdrd7* mRNA signal within a 3D germ granule that was segmented based on Vasa protein signal.
(B) Workflow of 3D germ granule layer analysis. The surface of large granules (0.8-2.5 µm radius) was segmented based on Vasa protein signal, and the minor axis length (radius, r) was determined. The upper right scheme shows a granule divided into layers with equal thickness (each ¼ of r), with the dotted outline representing the 'outer' layer that surrounds the granule surface (layer e). For the generation of 3D layers, the segmented granule surface was shrunk or expanded by a quarter of the granule radius (blue shaded box, surface in right image panels). Volumes were then generated based on the signal located within the different-sized surfaces (= masked signal). These volumes were subtracted from each other, generating 3D concentric layers (blue shaded box, shown as single planes in the left image panels). The layers were evaluated for integrity and subsequently the intensity and co-localization values were extracted for a particular RNA or protein type within each generated 3D layer. The intensity was normalized to the layer volume.
(C) Co-detection of endogenous *nanos3* mRNA and Dnd-GFP protein within germ granules. The image panels show an example of Dnd and *nanos3* localization within the condensate, with the white dotted outline highlighting the peripheral region of the granule. The diagram on the right panel shows the quantification of the overlap between the protein and RNA in the peripheral layer (layer d) of multiple granules (n = 21, N = 3).

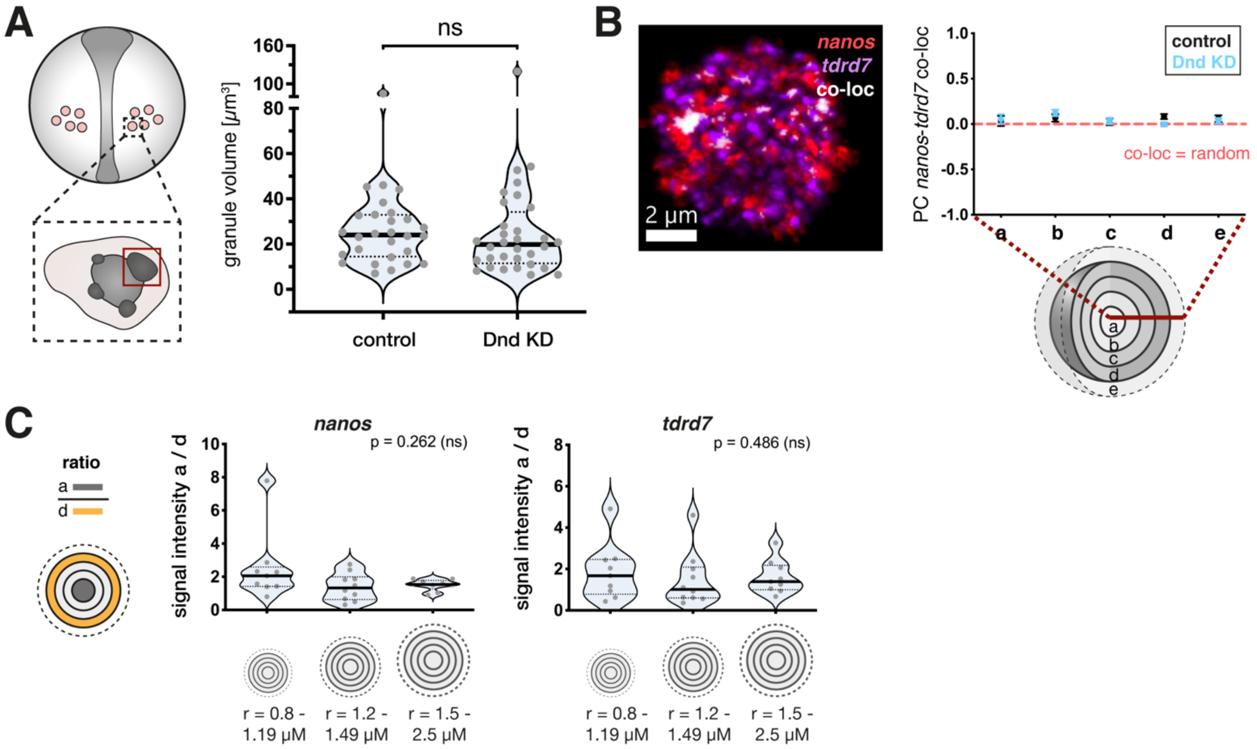


**Figure S2. Knockdown of Dnd does not affect germ granule volume nor homotypic clustering of germline mRNAs at 10 hpf, and different granule sizes do not affect mRNA organization, related to Figure 2.**

(A) Quantification of the volume of large condensates (red box in scheme) under control and Dnd KD conditions. Statistical analysis was performed using Mann-Whitney *U* test. ns = not significant. n (ctrl.) = 28, n (Dnd KD) = 32. Number of experiments performed (N) = 3.

(B) Evaluation of the co-localization between endogenous *nanos3* and *tdrd7* mRNA within the germ granule under control and Dnd KD conditions. Left image shows a 3D illustration of distinct mRNA clusters accumulating within a large germ granule condensate. The right graph shows the Pearson Coefficient (PC) of *nanos3* and *tdrd7* co-localization across the 3D germ granule layers. n (control) = 28, n (Dnd) = 32. N = 3.

(C) 3D analysis of the ratio of the mRNA level in the granule core (layer a) to that in the periphery (layer d) in large condensates that were categorized according to their size. r = radius. Significance was calculated using Kruskal-Wallis test. ns = not significant. n (r 0.8-1.19 µM) = 9, n (r 1.2-1.49 µM) = 10, n (r 1.5-2.5 µM) = 9. N = 3.

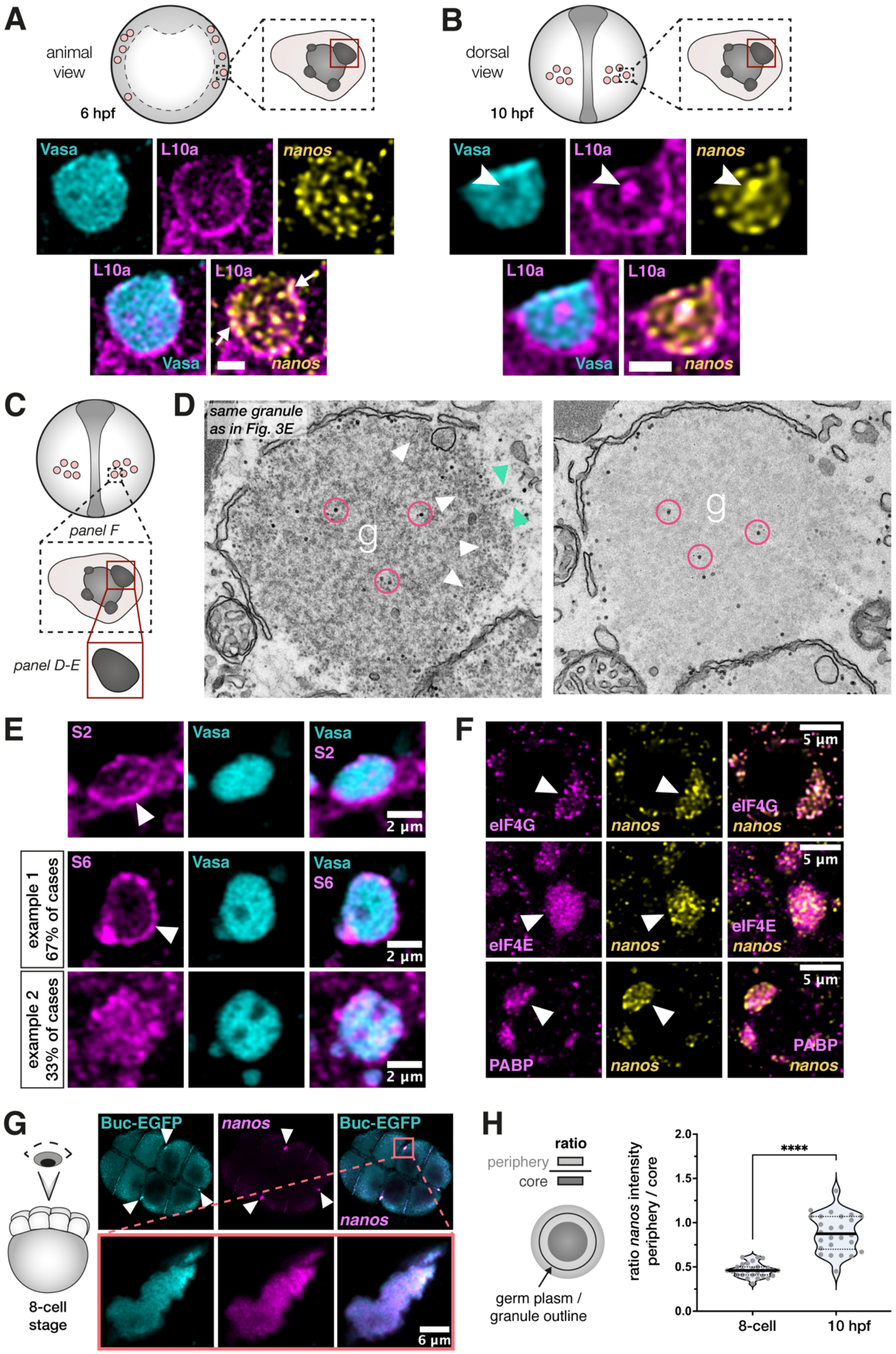

A
animal view
6 hpf
Vasa
L10a
nanos
L10a
Vasa
L10a
nanos
B
dorsal view
10 hpf
Vasa
L10a
nanos
L10a
Vasa
L10a
nanos
C
panel F
panel D-E
D
same granule as in Fig. 3E
g
g
E
S2
Vasa
Vasa S2
2 µm
example 1 67% of cases
S6
Vasa
Vasa S6
2 µm
example 2 33% of cases
2 µm
F
eIF4G
nanos
5 µm
eIF4G nanos
eIF4E
nanos
5 µm
eIF4E nanos
PABP
nanos
5 µm
PABP nanos
G
8-cell stage
Buc-EGFP
nanos
Buc-EGFP nanos
6 µm
H
ratio
periphery
core
germ plasm / granule outline
ratio nanos intensity periphery / core
2.0
1.5
1.0
0.5
0.0
****
8-cell
10 hpf

**Figure S3. Ribosomes are enriched at the periphery of zebrafish germ granule condensates, related to Figure 3.**

(A) Co-detection of Vasa protein, the 60S ribosomal protein L10a and endogenous *nanos3* mRNA in germ granules of 6 hpf embryos. Arrows point at overlaps between *nanos3* and L10a at the granule periphery. Scale bar = 2 μM.

(B) Co-detection of Vasa protein, L10a protein and endogenous *nanos3* mRNA in germ granules of 10 hpf embryos. Arrowheads point at an intra-granule spot of L10a and *nanos3* enrichment in a Vasa-deficient region. Scale bar = 2 μM.

(C) A scheme showing germ cells in a 10 hpf embryo. The black dotted box, and red box each depict the region of interest for the indicated image panels.

(D) Electron microscopy image of a germ granule condensate in a germ cell of a 8-12 hpf embryo, corresponding to Figure 3E. The two image panels show two consecutive ultrathin sections stained with uranyl acetate and lead (left panel) or lead only (right panel). In the left panel, free ribosomes (white arrowheads) and polyribosomes (green arrowheads) are enriched at the granule (g) periphery, and glycogen particles (magenta circles) can be detected. In the right panel, glycogen particles but not ribosomes are highlighted by the lead staining, allowing a clear distinction between the two structures.

(E) Co-detection of Vasa protein with 40S ribosomal proteins S2 (upper panels) and S6 (lower two panels). White arrowheads point to enrichment of ribosomal proteins at the germ granule periphery. Two examples of representative distributions patterns are shown for protein S6, which showed enrichment at the condensate periphery in most cases.

(F) Co-detection of endogenous *nanos3* mRNA and translation initiation factors eIF4G, eIF4E and PABP within germ cells. Arrowheads point at large germ granule condensates.

(G, H) Co-detection of endogenous *nanos3* mRNA and Bucky Ball-EGFP protein (Buc-EGFP) in 8-cell embryos. In the upper panels, an animal view on the embryo is presented, showing enrichment of germ plasm at the cleavage furrows of the blastomeres (white arrowheads). The red box marks a single germ plasm aggregate that is magnified in the lower panels. The graph compares the ratio of mRNA abundance between the granule periphery and the core of the germ plasm aggregate in 8-cell and 10 hpf embryos. Significance was calculated using Mann-Whitney *U* test. ****P<0.0001. n (8-cell) = 27, n (10 hpf) = 26. N = 3.

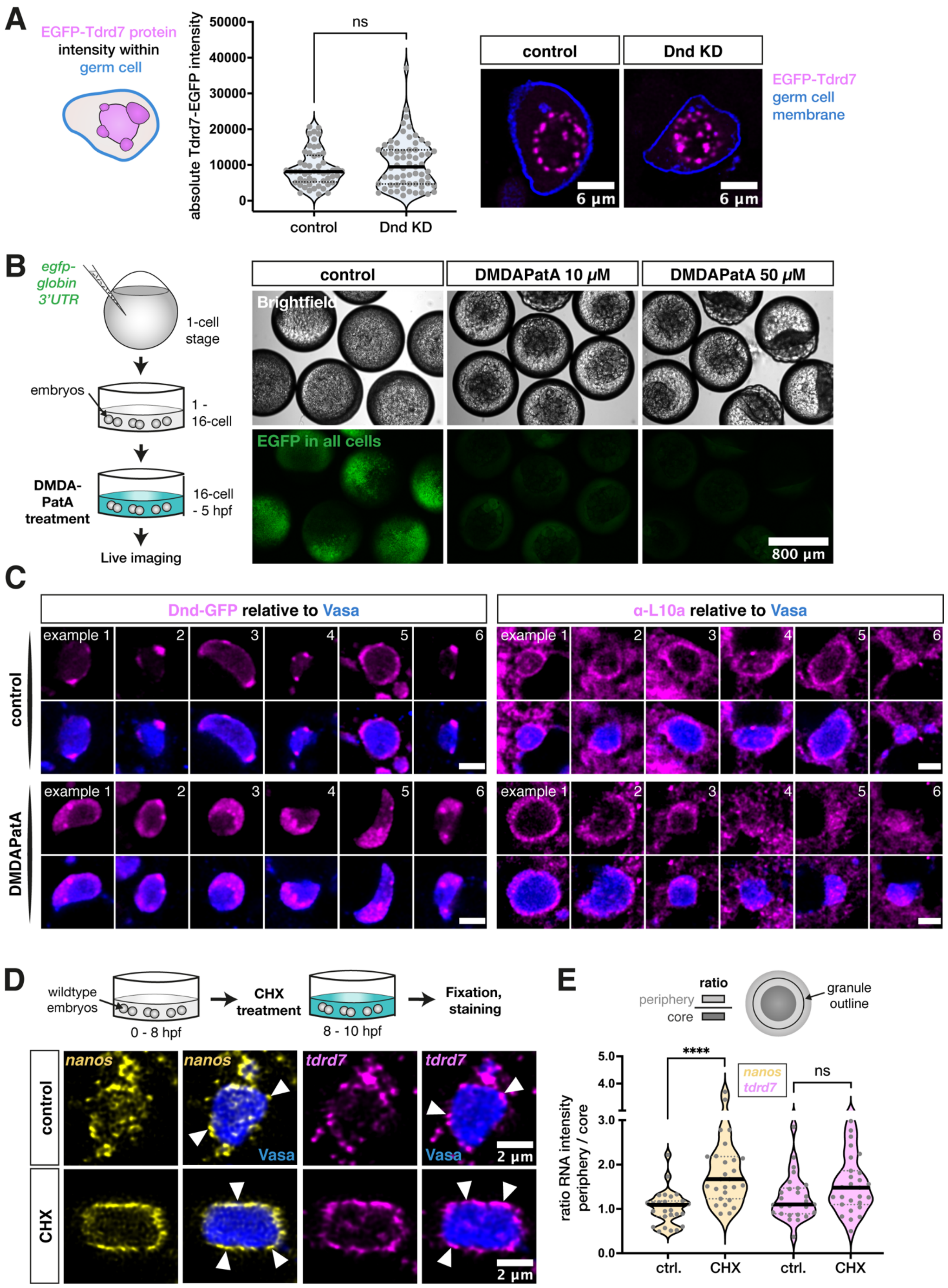
A
EGFP-Tdrd7 protein intensity within germ cell
absolute Tdrd7-EGFP intensity
ns
control
Dnd KD
EGFP-Tdrd7
germ cell membrane
6 µm
B
egfp-globin 3'UTR
1-cell stage
embryos
1 - 16-cell
DMDA-PatA treatment
16-cell - 5 hpf
Live imaging
control
DMDAPatA 10 µM
DMDAPatA 50 µM
Brightfield
EGFP in all cells
800 µm
C
Dnd-GFP relative to Vasa
α-L10a relative to Vasa
example 1
control
DMDAPatA
D
wildtype embryos
0 - 8 hpf
CHX treatment
8 - 10 hpf
Fixation, staining
nanos
tdrd7
Vasa
CHX
2 µm
E
ratio
periphery
core
granule outline
ratio RNA intensity periphery / core
****
ns
ctrl.
CHX

**Figure S4. *nanos3* mRNA localization to germ granule condensates depends on its accessibility to translation factors, related to Figure 4.**

(A) Evaluation of the mean signal intensity of EGFP-Tdrd7 protein, encoded by injected *tdrd7* 3'UTR-containing mRNA, within germ cells of control and Dnd KD embryos. The blue cell outline in the scheme represents the border of the area of measurement. Significance was calculated using Mann-Whitney *U* test. ns = not significant. n (ctrl.) = 36, n (Dnd KD) = 36. N = 3.

(B) Verification of DMDAPatA drug efficiency. 1-cell stage embryos were injected with mRNA encoding for globally expressed EGFP. The embryos were subjected to control (DMSO) or DMDAPatA treatment from the 16-cell stage to 5 hpf, followed by assessment of EGFP expression in live embryos.

(C) Localization of Dnd-GFP (left panels) and 60S ribosomal protein L10a (right panels) relative to Vasa protein in germ granules following control (upper panels) or DMDAPatA treatment (lower panels). For each protein, six representative examples are shown. Scale bar = 3 µM.

(D) Localization of endogenous *nanos3* mRNA and *tdrd7* mRNA relative to Vasa protein in germ granules following control or CHX treatment. Arrowheads point at clusters of mRNAs localized to the granule periphery.

(E) A graph showing the ratio of the mRNA level between the granule periphery and its core. Significance was calculated using Mann-Whitney U test. ****P<0.0001. ns = not significant. n (ctrl.) = 27, n (CHX) = 27. N = 3.

**Table S1. Oligonucleotides, related to STAR Methods.**
